# *Schistosoma mansoni* infection and risk factors among fishermen at Lake Hawassa, Southern Ethiopia

**DOI:** 10.1101/488502

**Authors:** Tadesse Menjetta, Daniel Dana, Serkadis Debalke

**Author notes:** (TM).

## Abstract

Schistosomiasis/Bilharziasis is one of the neglected tropical parasitic diseases caused by different species of genus schistosoma. Among the species, *S.mansoni* (causative agents of intestinal schistosomiasis) is one of the causes of severe intestinal parasitic infections with high public and medical importance in Ethiopia. There is scarcity of information about the status of *S.mansoni* infection among the fisherman in the present study area and in the country at large. Therefore this study was designed to determine the prevalence and risk factors of *S.mansoni* infection among fishermen at Lake Hawassa, southern Ethiopia. A cross-sectional study was conducted among the fishermen from April to June 2013 in Hawassa, Southern Ethiopia. A total of 243 fishermen were included by Systematic Random Sampling from the lists of the fishermen members in the registration book of fishermen associations in the Hawassa Town. Data on socio-demographic features and risk factors were collected by using semi-structured questionnaires. Stool samples were collected and processed using Kato-Katz thick smear techniques and examined between 30-40 minute for hook worm and after 24 hours for *S.mansoni* and other soil transmitted helminths (STHs). The overall prevalence of *S.mansoni* among the fishermen was 29.21% (71/243) and the mean intensity of infection was 158.88 eggs per gram (EPG). The prevalence of intestinal helminths including *S.mansoni* was 69.54% (169/243). Moreover, the prevalence of soil transmitted helminths (STHs) were 40.74% (99/243), 35.80% (87/243) and 5.76% (14/243) for *A. lumbricoides*, *T. trichiura* and hookworm species, respectively. Almost similar prevalence of *S.mansoni*, 31.82%, 31.75%, 31.94% were recorded in age groups of 15-19, 20-24 and 25-29 years, respectively. Fishermen who are swimming always were 2.92 times [95% CI: 1.554, 5.502] more likely to acquire *S.mansoni* infection than other water contacting habit of the study participants. The results of current investigation indicated the moderate endemicity of *S.mansoni* among the fishermen at Lake Hawassa, southern Ethiopia. Fishermen could be the potential risk group for *S.mansoni* infection and might be responsible for the transmission of *S.mansoni* to other segments of the communities. Since high prevalence of STH were recorded among the fishermen, integrated prevention and control strategies from different sectors might be important to tackle the problem.

**Author summary:** It is known that schistosomiasis is one of the neglected tropical parasitic diseases. However there is scarcity of information about the status of *S.mansoni* infection among the fisherman in the study area and in many parts of the world. Knowing the epidemiology of these parasites among the risk groups (fishermen) can contribute a lot to scale up the current control and elimination strategies. In addition, Fishermen could be the potential risk group for *S.mansoni* infection and might be responsible for the transmission of *S.mansoni* to other segments of the communities. To determine the prevalence of *S.mansoni* infection among the fisherman, the present study is done using stool samples from study groups.

## Introduction

Schistosomiasis is one of the chronic parasitic diseases caused by trematodes of the genus schistosoma. *S. mansoni* and *S. haematobium* are the major causes of human schistosomaisis in Ethiopia (1). *S.mansoni* and STH are the major causes of intestinal helminthiasis in developing countries and their impact on public health has been underestimated due to the chronic nature of the disease and low mortality rate (2). The distribution and prevalence of various species of intestinal helminths including *S.mansoni* differ from region to region due to environmental, social and geographical factors (3). In general, the burden of intestinal helminthiasis is high throughout the tropics, especially among poor communities and increasing trends of the infections recorded in developing nations including Ethiopia (4).

The global burden of schistosomiasis indicates about 200-209 million infected individuals and 600-779 million people at risk of infection (5). The estimated total numbers of people requiring treatment for schistosomiasis in 2014 were more than 258milion. About 91.4% of the people estimated to require treatment for schistosomiasis lived in the African Region (6). In Ethiopia, about 5.01 million people are thought to be infected with schistosomiasis and 37.5 million to be at risk (7).

Despite of the low mortality of *S.mansoni,* the chronic nature of the disease and the morbidity has an impact among fishermen, farmers, laborers who have frequent contact with contaminated water sources with *S.mansoni* cercaria (8). Among different risk factors for endemicity of *S.mansoni*, water conservation, irrigation, and hydroelectric power have contributed high for the spread of the disease and changed its epidemiology (9). Particularly, fishing, bathing, washing clothes and other activities involving frequent water contact could increase the transmission of *S.mansoni* (10).

Since the long-term aim of World Health Organization (WHO) is to eliminate schistosomiasis and STHs as a public health problem by 2020, the Federal Ministry of Health (FMOH), of Ethiopia has ambitious aims to scale up the treatment against all Neglected Tropical Diseases (NTDs), including schistosomiasis and STHs. Therefore, assessing the epidemiology of these helminths among the risk groups (fishermen) may contribute a lot to scale up the current control and elimination strategies.

From our observation, there are thousands of people around the present study area, Lake Hawassa, who are engaged in fishing and fish processing. Fishermen routinely swim, bath, and wash clothes in the Lake Hawassa. Those activities can easily expose fishermen for *S.mansoni* and other parasitic infections. Although different epidemiological studies of schistosomiasis have been undertaken among the school children in Ethiopia, the magnitude of *S.mansoni* and other intestinal helminths among the fishermen were not well addressed. Current study was undertaken to determine the magnitude of *S.mansoni* and risk factors among fishermen at Lake Hawassa. The findings may strengthen the information so far for scaling up the control and elimination measures of *S.mansoni* in the study area and in the country at large.

## Methods

### Study area and population

Hawassa Town is found in the southern part of Ethiopia, 250 km away from Addis Ababa, the capital of Ethiopia. Hawassa is the capital city of Southern Nations, Nationalities and People Region (SNNPR) with hot climate and has an altitude of 1680 m above sea level. The city is surrounded by Lake Hawassa which is part of east African rift valley belt. Lake Hawassa measures 16 km long and 9 km wide, with a surface area of 129 square kilometers and has a maximum depth of 10 meters with an elevation of 1,708 meters.

All people who are engaged in fishing activities at Lake Hawassa used as the source population and those systematically selected individuals from the fishermen association and who engaged in fishing activities during the study period were used as the study participants. Those fishermen who engaged in fishing for minimum of three months were included in the study while those fishermen who have taken any anti-schistosomal and antihelminthic treatment within the past three months were excluded from the study.

### Study design, sample size determination and sampling techniques

A cross-sectional study was conducted to determine the magnitude of *S. mansoni* and risk factors among fishermen at Lake Hawassa, Southern Ethiopia.

The sample size was calculated using the 95% confidence interval with 5% marginal error.

Using the formula 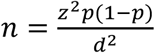, where n= sample size, z = z statistic for a level of confidence (z =1.96 at 95% CI), p = prevalence (p = 0.50), d = precision (if 5%, d= 0.05), the sample size became 384. Since sampling is from finite population of less than 10,000, population correction was used.

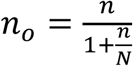; Where n= the sample size from the finite population, N= the total number of study population, and *n*_*o*_= the corrected sample size. Then 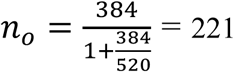. Allowing 10% for non response, the final sample size became 243.

Systematic Random Sampling technique was used to select study participants from the sample frame of the lists of fishermen in the fishermen association registration book. The following steps were followed: The interval size, k = N/n = 520/243= 2.13∼ 2. After the random selection of the 1^st^ participant, every 2^nd^ unit from the fishermen was included until the 243 fishermen were included.

### Stool sample collection and examination

Before the stool sample collection, semi-structured questionnaire was used to collect socio demographic data and risk factors by trained data collectors. The study participants were oriented how to provide sufficient stool specimen by principal investigator. A fresh faecal specimen was collected using labeled, clean, dry and leak-proof container. The stool specimen was processed using Kato-Katz thick smear technique. Each Kato-Katz smear was read with 30-40 minute for the detection and identification of hookworms and again re-read after 24 hours for *S.mansoni* and other helminths. The number of eggs of each species was recorded and converted into the number of EPG of stool in order to analyze intensity of infection. EPG in Kato-katz was calculated by multiplying the egg count by conversion factor that is 24 (11).

Intensity of infection was graded as light, moderate and heavy by counting helminths eggs excreted in faeces. The intensity for *S. mansion* is considered as light: 1-99 EPG, moderate: 100-399 EPG and heavy: ≥ 400 EPG. The intensity for *T trichiura* considered as light: 1-999 EPG, moderate: 1000-9999 EPG, heavy: ≥10,000 EPG. The intensity for *A.lumbricoides* considered as: light: 1-4999EPG, moderate: 5,000-49,999 EPG, heavy: ≥50,000 EPG and the intensity for hookworm considered as light: 1–1,999 EPG, moderate: 2,000–3,999 EPG, heavy: >3,999 EPG (12).

Standard operating procedures (SOP) were followed during stool specimen collection, transportation, processing, examination and result recording. From all positive and negative Kato-Katz thick smears, 10% were randomly selected and read by experienced laboratory technologist who is blind to the primary result and agreement was made for discrepancies.

### Data analysis

All questionnaires for socio-demographic and risk factors were checked for completeness. The data was coded, entered, cleaned and analyzed using SPSS version 20.0 statistical packages. Those variables with significant association in bivariate analysis were fitted to multivariate analysis to determine the independent predictors and statistical significance was considered as value < 0.05.

### Ethical consideration

Ethical clearance was obtained from the Institutional Review Board (IRB) of Jimma University, College of public health and medical sciences. After explaining the objective and the purpose of the study, written informed consent was obtained from each fisherman involved in the study. Confidentiality of the information was assured and privacy of the respondent was maintained during the study. Results of participants were kept confidential and fishermen infected *S.mansoni* and other intestinal helminths were treated with praziquantel of 40mg/kg body weight and albendazole of 400mg according the WHO guideline by experienced health professionals.

## Results

### Socio-demographic characteristics

A total of 243 fishermen with age range of 15 to 39 years were participated in this study. Majorities of the study participants were between 25-29 years age group. About 73.25% (178/243) of the study participants were in elementary level of education. About 58.44% (142/243) were from urban and 41.56% (101/243) were from rural areas (Table 1).

### Prevalence of *S.mansoni* and other intestinal helminthic infection

The prevalence of *S.mansoni* among the fishermen was 29.22 % (71/243).The prevalence of 31.82%(14/44), 31.75%(20/63), 31.94%(23/72),17.07%(7/41) and 30.43%(7/23) were recorded among the age groups of 15<19, 20<24, 25<29, 30<34, and 35<39 years, respectively. Of the total 243 stool samples examined, 169 were positive for one or more of intestinal helminths giving an overall prevalence of intestinal helminths to be 69.54%. The prevalence of *A. lumbricoides, T. trichiura* and hookworm species were 40.74 % (99/243), 35.80 % (87/243), and 5.76 % (14/243), respectively (Figure 1).

### Multiple intestinal helminthic infections

Of 169 helminth infected fishermen, 43.78% (74/169) of them had single helminths infection, while 56.21% (95/169) were infected with more than one intestinal helminth. The most common multiple helminths infection were double infection of *A. lumbricoides* and *T. trichiura (*50.52 %) followed by *S.mansoni* and *T.trichuria* (13.68%).The major triple infection (9.47%) were *S.mansoni, A. lumbricoides* and hookworm species (Table 2).

### Intensities of S.mansoni and Soil transmitted helminth

The intensities of *S.mansoni* and STHs have been assessed using faecal egg count by Kato-Katz thick smear. The mean intensity of *S.mansoni* infection was 158.88 EPG. The level of infection intensities for *S. mansoni* among the fishermen were 52.11%, 43.66% and 4.23% for light, moderate and heavy infection intensities, respectively. The mean infection intensity of *A.lumbricoides, T. trichiura* and hookworm were 1349.04 EPG, 246.24 EPG and 99.36 EPG, respectively. Light infection intensity of *A.lumbricoides* and *T.trichiura* were recorded in 81.82% and 91.95% of the fishermen, respectively. Only light infection intensity was recorded in hookworm infected fishermen (Table 3).

### Risk factors associated with S. mansoni infection

*S. mansoni* infection among fishermen was significantly associated (P<0.05) with swimming, bathing and other activities with frequent water contact. On the other hand, there is no statistically significant association between *S. mansoni* infection and water sources utilized by the fishermen. Swimming, frequency of swimming and frequency of water contact were the independent predictors for *S.mansoni* infection among the fisherman. Fishermen who used to swim always in Lake Hawassa were 2.92 times [95% CI: 1.554, 5.502] more likely to acquire *S. mansoni* infection than those who swim some times. But frequency of water contact and habit of swimming were not a risk even if they have association in the bivariate analysis. However there is no statistically significant association between *S. mansoni* infection and the place where the fishermen bath (Table 4).

## DISCUSSION

Schistosomiasis is one of the neglected tropical parasitic diseases with multiple risk factors related to the parasite, environment, human and snails (13). Based on the nature of occupation, fishermen could be easily infected by schistosomiasis and might be a potential source for the transmission the disease to other segments of population. The overall prevalence of *S. mansoni* among fishermen at Lake Hawassa in the present study is found to be 29.22%. The majorities (52.11%) of infection intensities were low level infection intensity. The prevalence of *S.mansoni* among the fishermen in East Africa are higher than the prevalence in Lake Hawassa, including the prevalence of 88.6% (14), 72% (15) and 47.4% in Uganda (16), 47.85% in Tanzania (17), 72.4% in Egypt (18) and 41.3% in Ethiopia (19). The difference of prevalence might be due to the variation in geography, climate and ecology, the study design, awareness level and immune status of the fishermen. Moreover, Uganda and Tanzania are among the world’s top fishing nations and there are large segment of population engaged fishing than in Ethiopia. Similarly, higher prevalence of *S.mansoni* (33%) were reported in the same study area (Lake Hawassa, Ethiopia) among the children engaged in fishing and fish processing activities (20). The variation of prevalence might be due to the frequent playing and swimming habit of the children in Lake Hawassa.

The prevalence finding of our study area is higher than some of the prevalence reports among the fishermen from Burkina Faso (16.35%) (21) and Egypt (26.6%) (22). Similarly, lower prevalence (12.5% of *S.mansoni* was reported among subsistence fishermen and commercial fishermen in Zambia (23).The difference in the prevalence might be due to geographic, climatic and ecological factors. Moreover, the personal and environmental sanitation level might be responsible for the difference of *S.mansoni* prevalence from place to place.

According to different literatures, there is high chance of co-endemicity for schistosomiasis and STHs in areas where schistosomiasis is prevalent (24). Even though it is not our primary objective, we assessed the prevalence of other intestinal helminths in our study. The overall prevalence of intestinal helminths including *S.mansoni* among fishermen was 69.55 %. Among those intestinal helminths, STHs are the major helminths in the fishermen in our study area. The prevalence of STHs were 40.74%, 35.80% and 5.74% for *A. lumbricoides, T. trichiura* and hook worms, respectively. The prevalence of *A. lumbricoides* is comparable with the prevalence finding among fishermen in Vietnam, but higher prevalence of *T. trichiura* and hook worm was reported from Vietnam (25). Similarly, higher prevalence of all the three STHs (*A. lumbricoides, T. trichiura* and hook worms) were reported among children in fishing villages in India (26). The difference in prevalence might be due to geographic, climatic, socioeconomic status, environmental and personal hygiene level of study participants. In the contrary, the prevalence of the three STHs in our study area is higher than the results reported among the community members mainly engaged in fishing in rural Abaya Deneba area that is adjacent to Lake Ziway, Ethiopia (19). The variation of prevalence might be due to the difference in the diagnostic methods employed.

The present study is subject to some limitations. Since the study design was cross-sectional; it is difficult to make causal inference and to study the transmission dynamics of *S.mansoni* infection. Due to the deficit of literature regarding the epidemiology of *S.mansoni* and risk factors among fishermen, it was difficult to discuss the risk factors. Moreover, those few studies of *S.mansoni* among fishermen are not reporting other helminthic co-infection like the commonly diagnosed STHs in kato-katz techniques.

## Conclusions

The results of current investigation indicated the evidence for moderate endemicity of *S.mansoni* among the fishermen at Lake Hawassa, southern Ethiopia. Fishermen could be the potential risk group for *S.mansoni* infection and might be responsible for the transmission of the disease to other segments of the communities. Unexpectedly, high prevalence of STH was recorded among the fishermen. Therefore, integrated prevention and control strategies for both schistosomiasis and STHs from different sectors might be important to tackle the problem.

## Acknowledgements

The authors would like to thank the Hawassa College of Health Sciences, for provision laboratory facility for the examination of stool samples. Our deepest gratitude also goes to the Hawassa Fishermen Association for their cooperation during data collection.

## Data Availability

All relevant data are within the paper.

## Abbreviations

FMOH: Federal Ministry of Health
SNNPR: southern nation and nationalities people region
STHs: soil transmitted helminths
EPG: egg per gram
COR: Crude Odds Ratio
CI: Confidence Interval
SOP: Standard operating procedures

## Author information

TM, MSc in medical parasitology from Jimma University, Ethiopia, working at Hawassa University, College of Medicine and Health Science, School of Medical Laboratory Sciences, Hawassa, Ethiopia.

SD, MSc in infectious disease from Addis Ababa University, Ethiopia, PhD fellow in Ghent University, Belgium.

DD, MSC in Medical Parasitology from Jimma University, Ethiopia, PhD fellow in Ghent University, Belgium.

## Supporting Information Legends

S1 Table. Socio-demographic characteristics of the fishermen at Lake Hawassa, Southern Ethiopia, 2013

S2 Table. Pattern of multiple intestinal helminthic infections among fishermen at Lake Hawassa, 2013

S3 Table. Intensity of *S. mansoni, A. lumbricoides, T. trichiura* and hookworm infections using Kato-Katz thick smear technique among fishermen at Lake Hawassa, 2013

S4 Table. Bivariate and Multivariate logistic regression analysis of factors associated with *S. mansoni* infections among fishermen at Lake Hawassa, Southern Ethiopia, 2013

S6 Fig. Prevalence and species distribution of intestinal helminth among the fishermen at Lake Hawasssa, 2013

S7 Checklist: STROBE Checklist

